# Periodicity in the embryo: emergence of order in space, diffusion of order in time

**DOI:** 10.1101/2021.01.06.425581

**Authors:** Bradly Alicea, Jesse Parent, Ujjwal Singh

## Abstract

Does embryonic development exhibit characteristic temporal features? This is apparent in evolution, where evolutionary change has been shown to occur in bursts of activity. Using two animal models (Nematode, *Caenorhabditis elegans* and Zebrafish, *Danio rerio*) and simulated data, we demonstrate that temporal heterogeneity exists in embryogenesis at the cellular level, and may have functional consequences. Cell proliferation and division from cell tracking data is subject to analysis to characterize specific features in each model species. Simulated data is then used to understand what role this variation might play in producing phenotypic variation in the adult phenotype. This goes beyond a molecular characterization of developmental regulation to provide a quantitative result at the phenotypic scale of complexity.

## Introduction

While the case for the effects of “tempo and mode” [1] have been made for the evolutionary process, a similar relationship between phenotypic change, time, and space may also exist in development. One obvious answer to this question is to examine the expression and sequence variation of genes associated with cell cycle and developmental patterning [2]. However, there is a potentially more compelling top-down explanation. We will use two model organisms to demonstrate how periodicity becomes less synchronized over developmental time and space. In the case of the nematode *Caenorhabditis elegans*, a comparison of embryogenetic and postembryonic cells (developmental and terminally-differentiated cell division times acquired from [3]) reveals two general patterns. For the Zebrafish (*Danio rerio*), comparisons within and between embryogenesis stages based on measurements of cell nuclei in the animal hemisphere [4] reveal patterns at multiple scales. One of the most notable signatures is burstiness [5, 6], or a large number of events occurring in a short period of time. These bursts can either be periodic or aperiodic, and these statistical features define the temporal nature of development, potentially in a universal manner across species. An example of burstiness and other temporal dynamics during the embryonic development of Zebrafish can be observed in Supplemental Movie 1 [7].

Based on two species and a computational model, we predict that periodic changes in the frequency of new cells over developmental time will exhibit stochastic variation not necessarily related to differentiation into specific cell fates [8]. We also analyze the intervals between bursts in cell division (and cell differentiation in the case of *C. elegans*). This bursty behavior [9] is derived from both time-series segmentation and decomposition in the frequency domain. We consistently observe great temporal variation at the cellular level, and may play a role in shaping morphogenesis. In addition, these changes in frequency and periodicity over time produce spatial variation (Supplemental Figure 1). To characterize spatial variation, we utilize embryo networks [10]. Embryo networks are complex networks based on the relative proximity of cells as they divide and migrate during the developmental process. The resulting network topologies provide not only information about spatial variation, but cellular interactions and other signaling connections as well [11, 12]. The existence of network structure in the form of modules or regions of dense connectivity can reveal a great deal about the unfolding of lineage trees in time.

Returning to the first prediction, we can create computational summaries of cell division events called numeric embryos to model the proliferation of cells over time. We call these computational models, numeric embryos, and can be used to model branching events in a lineage tree. Numeric embryos can be used to model the distribution of branching events in time, independent of cell identity or spatial context. Approximating this distribution provides us with a periodic time-series that tells us something about the speed of embryogenesis: how quickly can different underlying distributions of cell division produce a phenotype with many developmental cells (cells yet to reach terminal differentiation). The rate at which developmental cells are produced could affect the rate of overall development, as we will see in an example from Zebrafish.

Finally, we predict that the emergence and subsequent changes in spatiotemporal periodicity at the cellular level lead to regulatory phase transitions. For example, there is a one-to-one correspondence between cell division and waves of differentiation after the syncytial stage in *Drosophila melanogaster* [13]. In a similar fashion, amphibians exhibit a decay of synchrony of division [14, 15] that might correspond to the unfolding of lineage and differentiation trees [16]. Differentiation trees serve as the basis for differentiation waves [17], which further the link between temporal and spatial phenomena. Based on data analysis, modeling, and literature review, we anticipate that further investigation could uncover whether, in regulating embryos, mitosis and cell differentiation are correlated. In interpreting the data, we discuss the potential applicability of Holtzer’s quantal mitosis hypothesis [18, 19] as it relates to the process of differentiation relative to the proliferation of cells yet to reach terminal differentiation.

## Methods

A summary of the methods could be given here for smooth reading and interest. All materials are located on Github [20]. This repository includes processed data, supplemental materials, and associated code.

### Secondary Datasets

The *C. elegans* and *D. rerio* data sets were acquired from the Systems Science of Biology Database [21]. The *C. elegans* (nematode) data is based on cell tracking of nuclei [see 22]. The *D. rerio* (Zebrafish) data is likewise based on cell track of nuclei [see 23]. The cell tracking data is used to determine the total number of new cells (cell division time) present at a particular time step.

For the *C. elegans* data, cell divisions correspond to minutes of developmental time, and windows of size five (5 minutes of developmental time) is used for the time-series plots and histograms. Since lineage trees and the nature of developmental cell identification are different in Zebrafish, cell births correspond to the number of observed cells at discrete points in developmental time. Windows representing a certain number of cells in the embryo observed at a given sampling point are used instead of directly converting this process to minutes of developmental time.

### Zebrafish Developmental Stages

Estimates and calculations of *D. rerio* developmental stages are derived from [24] and the ZFIN Zebrafish Developmental Staging Series web resource (https://zfin.org/zf_info/zfbook/stages/). Where applicable, embryo stages are approximated from the number of cells observed at any given point in developmental time.

### Peak-finding method

For both the *C. elegans* and *D. rerio* data, a peak finding method is used to evaluate periodicity and to generate data points representing distinct bursts of cell divisions. Briefly, temporally local peaks in the cell division series are discovered by finding the highest value around the peak over an interval of 10 data points. The data are then visually inspected to ensure that temporally local (as opposed to temporally global) maximum fluctuations were not selected. Using this segmentation method, we are able to define intervals between peaks in a way that allows for the aperiodic regions of our series to be compared to the highly periodic regions.

The peak finding method result is supplemented by a Fast Frequency Analysis (FFT) of cell divisions in *C. elegans* embryo (Supplemental Figure 2), cell differentiation events in *C. elegans* embryo (Supplemental Figure 3), and time series for cell divisions in Zebrafish embryo (Supplemental Figure 4). The power spectra largely confirm the nature of our interval and peak analysis. While the analysis of Zebrafish reveals a power spectrum at a single scale, the *C. elegans* embryo reveals a power spectrum of multiple time scales for both cell divisions and differentiations.

### Embryo Networks

The full methodology for constructing and evaluating can be found in [10]. Briefly, embryo networks are complex networks constructed from the locations of cells in an embryo. Nodes are represented by centroids representing cell nuclei, and edges represent the spatial (Euclidean) distance between cells in a three-(static) or four-(dynamic) dimensional graph. All nuclei are plotted in embryo space, which is a coordinate system normalized to the center point between all cell locations in a complete embryo. For example, an edge of length 1.0 represents two centroids at opposite edges of the embryo space. A distance threshold is then derived from the length of the edge: in this paper, a distance threshold of 0.05 is used, excluding all but the cell nuclei in very close proximity to each other.

### Numeric Embryo

Numeric embryos are statistical summaries of the type of information acquired from our secondary datasets, but in a more generic manner. Numeric embryos are based on generated pseudo data and are meant to capture the structure of hypothetical developmental scenarios. All analyses of our pseudo data were conducted using SciLab 6.1 (Paris, France). Each numeric embryo consists of one or more vectors describing rounds of cell division in the embryo. Briefly, each minute of developmental time is represented by either a zero or a positive non-zero value. For purposes of temporal comparison, all non-zero values are thresholded to one. To generate cell division intervals of different sizes, we start with a uniform distribution (division events occur every *n* minutes) and then compare this with a distribution generated using the grand function in SciLab. For the Poisson distribution, we use a λ = 0.1 (except where otherwise noted), while for the Binomial distribution, we use parameters *N* = 1.0 and *p* = 0.5. This produces intervals that are variable over developmental time.

## Results

Our analysis will proceed from *C. elegans* to Zebrafish, to a comparison of the two species, then to a network analysis, and finally to a simulation of cell division in development. First, we plot the developmental cell division dynamics in *C. elegans* and Zebrafish in Figures 1 and 3, respectively, and cell differentiation in *C. elegans* in Figure 1. We then examine the intervals between cell division events (*C. elegans*) and relative frequency of birth rates across development (Zebrafish) in Figures 2 and 4, respectively.

**Figure 1.**
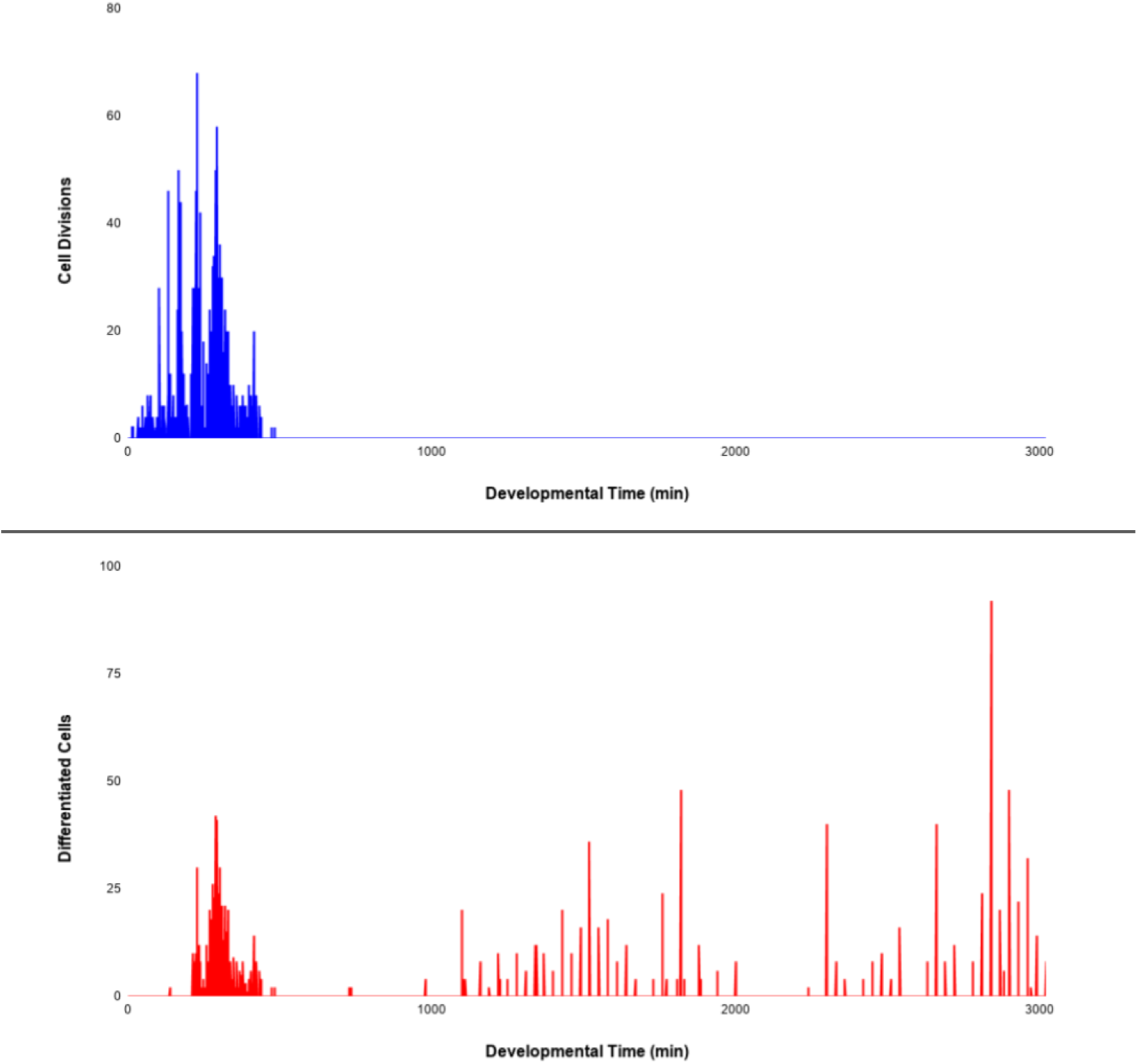
Developmental cell divisions in the nematode *C. elegans*. Cell divisions occur according to developmental time (minutes). The timeline ranges from fertilized egg (zygote) to adulthood. Embryonic division events (blue), terminal differentiation events to adult cells (red).

**Figure 2.**
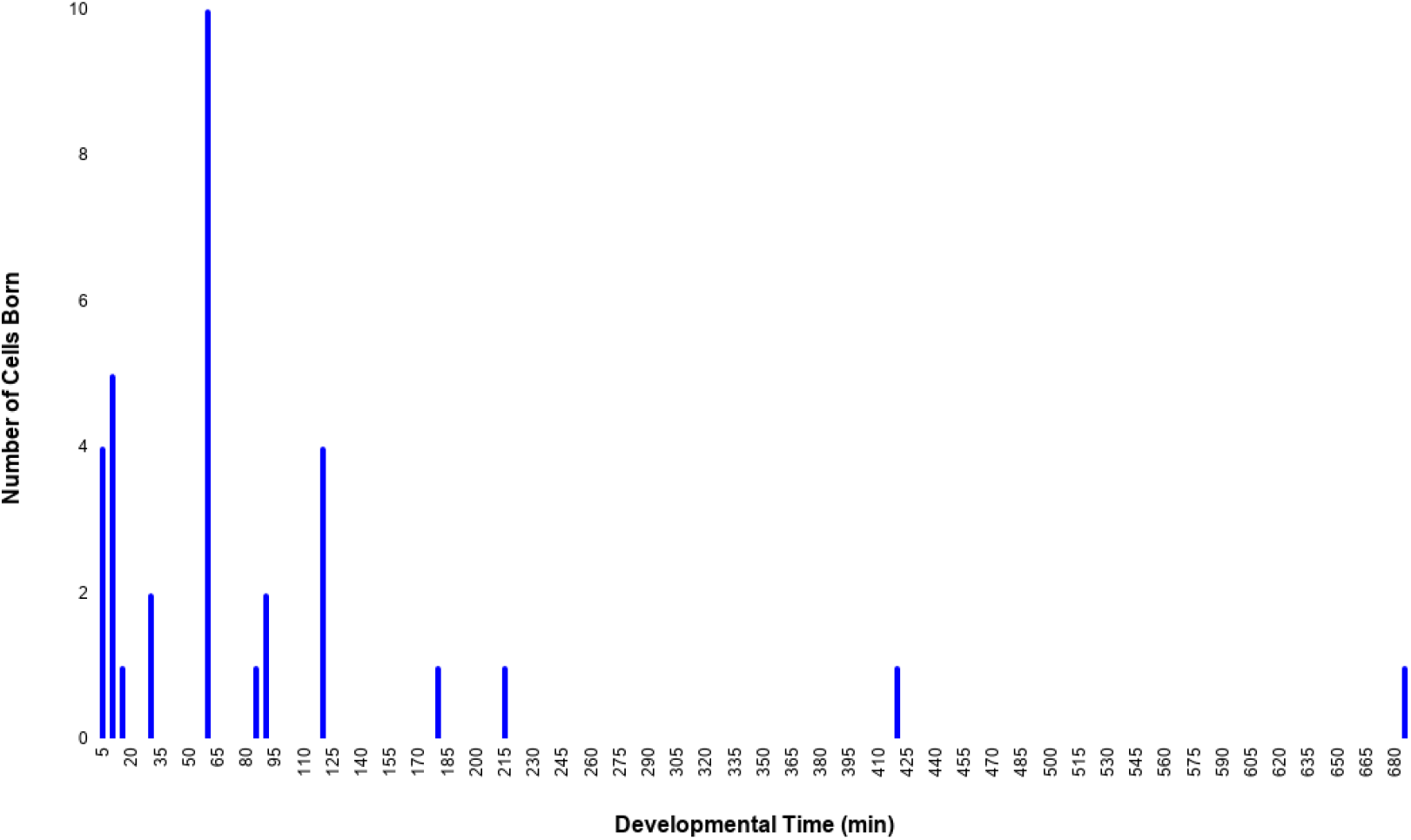
The interval between cell division events across embryonic development in *C. elegans*.

**Figure 3.**
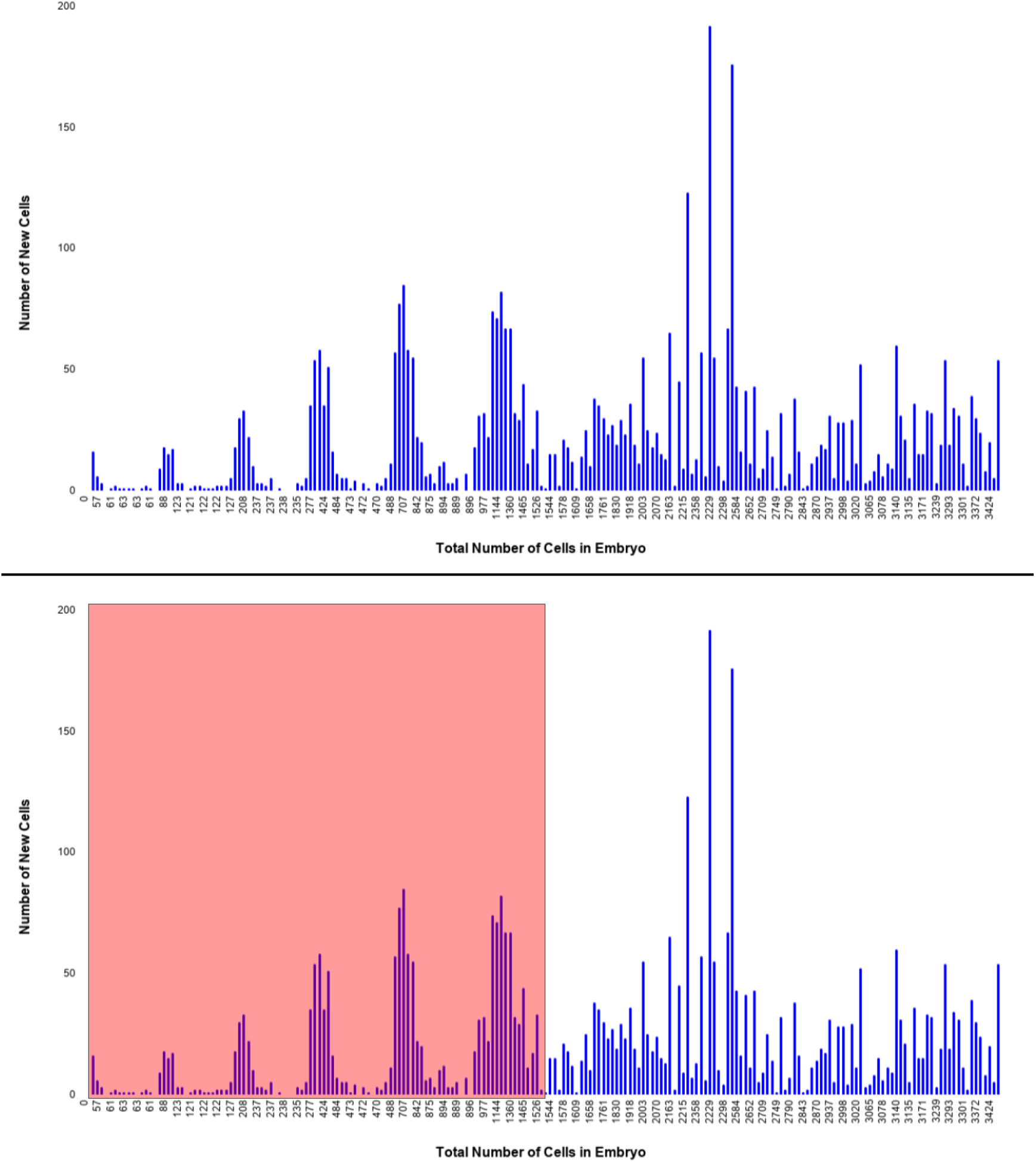
Cell divisions in Zebrafish embryos during embryogenesis up to the Gastrula stage. Instead of developmental time, relative developmental progress is plotted as all cells observed in the embryo at each sampling time point. For Figure 3, bottom: Periodic region (red), Aperiodic region (unshaded).

**Figure 4.**
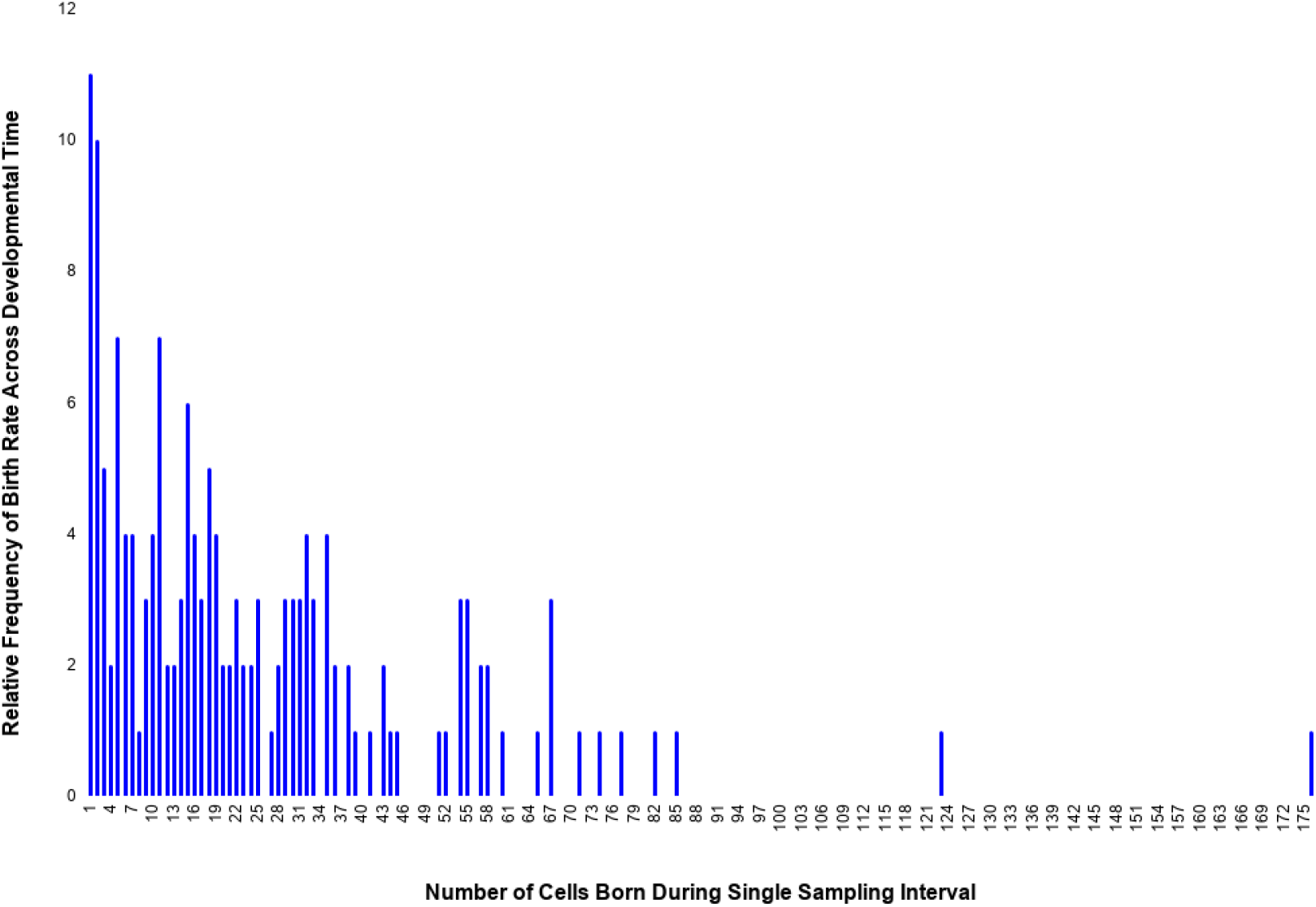
Relative frequency of division rate across developmental time in *D. rerio*. Histogram demonstrates the distribution of cells born during a single sampling time point.

Focusing on the peaks (maximum of bursts of cell divisions) shown in Figures 1 and 3, Figure 5 shows the distribution of intervals between peak values for *C. elegans* and Zebrafish. Figure 6 helps us extend this finding from temporal dynamics to connectivity between cells and spatial distributions of newly-born cells. We conclude with an investigation of how the intervals found between cell divisions can be modeled using various statistical distributions and is shown in Figure 7. These simulations (called numeric embryos) can reveal properties related to the speed of development, particularly the linear and nonlinear accumulation of cells.

**Figure 5.**
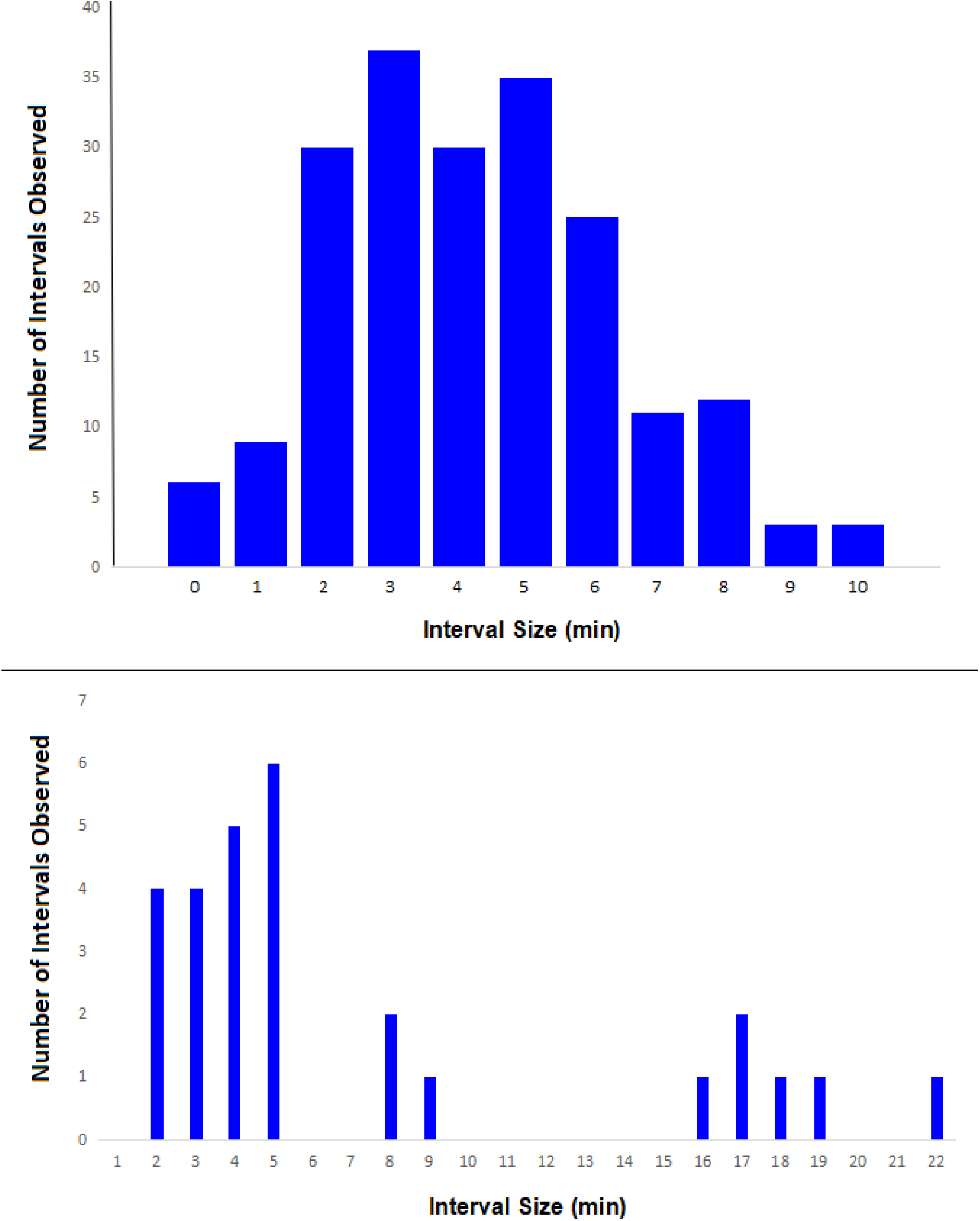
Interval size of peaks in cell division for all developmental cells in *C. elegans* (top) and first 206 minutes of Zebrafish (bottom). *C. elegans* sampling time points correspond to most of the pre-hatch developmental period (660 minutes post-fertilization), while the Zebrafish sampling time points correspond roughly to the period between the Zygote and the oblong/sphere stages of the Blastula.

**Figure 6.**
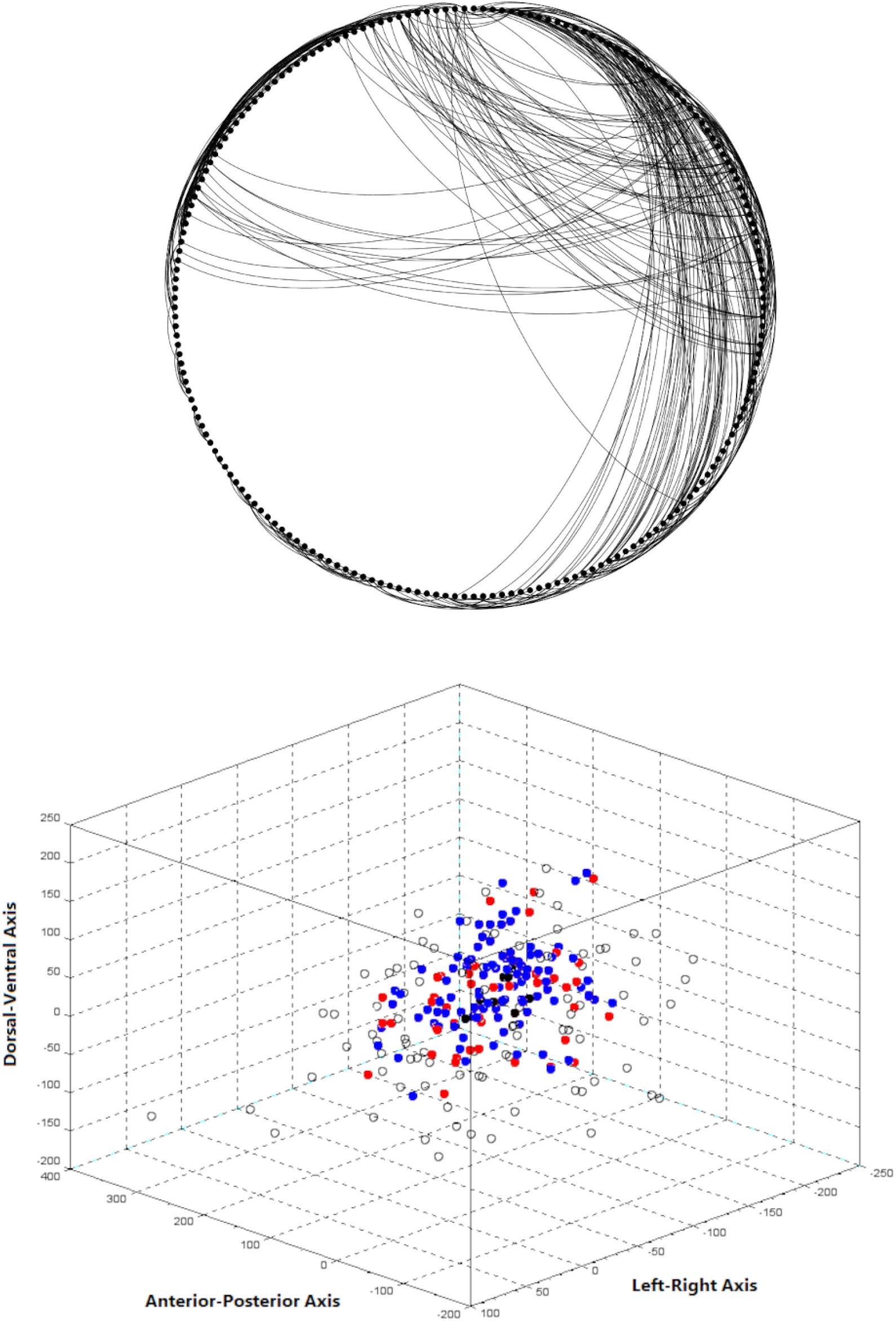
TOP: an embryo network for the *D. rerio* embryo at the 239 cell stage (all cells born during the zygote and cleavage stages), with 920 edges. The edge threshold is an embryo distance of 0.05. BOTTOM: Cells in developmental location color-coded by status in the network. WHITE: all cells not above the threshold, RED: all source cells with at least one edge to another cell. BLUE: all destination cells with at least one edge to another cell. Red and blue are equivocal. BLACK: all cells with more than eight edges to other cells.

**Figure 7.**
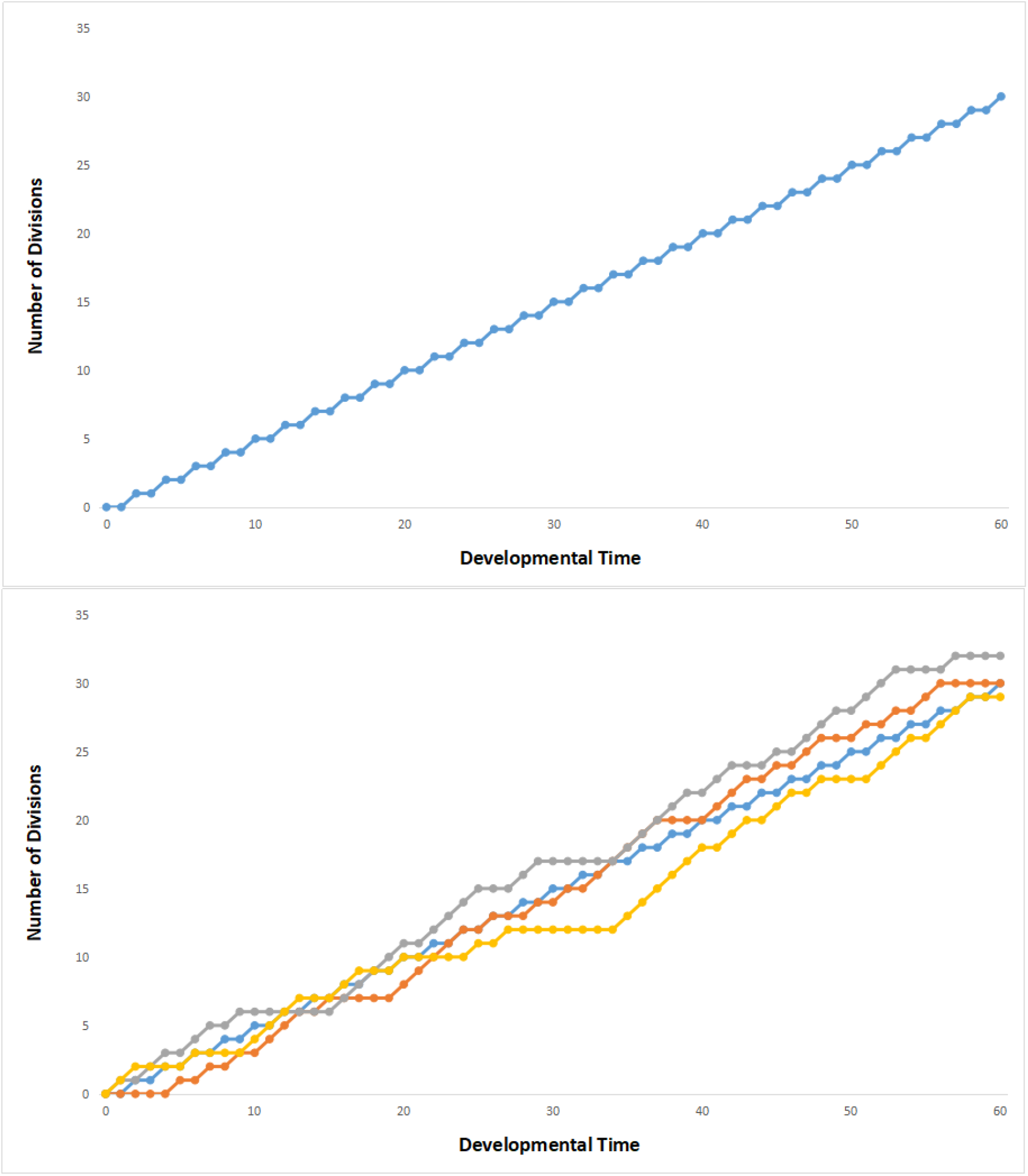
Comparison of cumulative cell division events and the speed of division generated by a numeric embryo. Top: Uniform only (blue). Bottom: Uniform (blue), Exponential (orange), Poisson (gray), and Binomial (yellow).

### *Caenorhabditis elegans* Example

To understand the temporal nature of cell division and differentiation, we start by looking at patterns in *C. elegans* development over time. Figure 1 shows a time series of such events from zygote to adulthood. We are particularly interested in potential spikes or bursts of events in a short period of time.

Figure 1 shows the fluctuations in cell divisions in embryonic division (Figure 1, top) and terminal differentiation events (Figure 1, bottom) events. Differentiation events occurring after 1000 minutes of developmental time (postembryonic development) occur in a long series of bursts, likely corresponding to the production of new seam cells. Seam cells are defined as lateral hypodermal cells that cover the seam on the medial sides of the worm. Seam cells also divide post-embryonically, before each larval molt [25]. This can be contrasted with the burstiness that occurs in embryonic development, which is similar to the burstiness of division events.

Figure 2 shows the intervals between cell division events across embryonic development in *C. elegans*. This plot confirms an exponential distribution with a long tail, presumably representing intervals in postembryonic development. Yet this plot is also sparse, yielding only 12 distinct intervals of cell division throughout all of *C. elegans* development. This is likely due to the deterministic nature of *C. elegans* development along with the relatively small number of cells. Supplemental Figures 2 and 3 reveal the power spectrum for cell division and cell differentiation in *C. elegans*, respectively. To compare, contrast, and understand these trends further, we now turn to the embryonic development of the Zebrafish (*D. rerio*).

### Zebrafish

In Figure 3 (top), we observe six regular busts of cell division, followed by aperiodic cell division behavior. This transition in periodicity is observed after the embryo reaches 1529 cells in size (Figure 3, bottom). We do not observe this in *C. elegans* embryos, and may have to do with the more regulative nature of Zebrafish embryogenesis [26]. Changes in periodicity may also have to do with the establishment of spatial differentiation beyond the axial variability observed in *C. elegans*.

To better understand the nature of periodicity in Zebrafish, we examined the distribution of intervals between birth times. Figure 4 and Supplemental Figure 4 confirms the bursty nature of cell division in Zebrafish, in that most sampling time points only feature a few cell births, while a small number of sampling time points represents a large number of cells born. For example, a large majority of sampling time points feature fewer than 25 new cells per time point. By contrast, there are also single sampling points where over 70 cells are born at a single time. In terms of the power spectrum shown in Supplemental Figure 4, there is a very high amplitude at very low frequencies, perhaps related to the significant noise and aperiodicity in the later part of the time-series shown in Figure 2.

Considering the cell divisions for the first period of Zebrafish embryogenesis, we conduct an interval analysis for each oscillation of the data shown in Figure 5 for *C. elegans* (top) and *D. rerio* (bottom). These are measured from peak to peak as described in the Methods. For the analysis of *C. elegans* data (Figure 5, top), our analysis yields a roughly unimodal distribution, with a mean peak interval of 3-5 minutes. In pre-hatch *C. elegans* embryogenesis, there are many quick bursts of cell division as confirmed in Figure 1 (top). This results in bursty behavior that is regular and perhaps even periodic.

By contrast., an analysis of our Zebrafish data yields three interval groups (Figure 5, bottom): the greatest number of oscillations occurs at a period of 2-5 minutes, while a smaller number of oscillations occur with periods from 16-19. There is also a longer 22-minute interval between oscillations. This is consistent with the shift from periodic bursts to aperiodic but still bursty behavior later in Zebrafish development shown in Figure 3. This multimodal distribution of peaks points to a more complex process at play, something that might be better understood by investigating morphogenesis as a spatial process.

### Embryo Networks: an example from Zebrafish

Another way to identify the consequences of bursts in cell division timing and other non-uniform temporal phenomena is to utilize embryo networks. An embryo network was constructed (Figure 6, Top) for cells born during our sampling time points of *D. rerio* embryogenesis. The resulting circular graph demonstrates a high degree of modularity, but only across part of the graph.

A three-dimensional plot (Figure 6, Bottom) demonstrating the position of each cell born during these stages of development shows that the highest degrees of connectivity are clustered in the center of the embryo, while cells that are disconnected based on our connectivity threshold exist on the edges of the embryo. Importantly, it appears that cells are more densely clustered toward the center of the embryo early at the earliest stages of development. These dense clusters are likely the product of cell division fluctuations shown in Figures 3 and 4.

One quantitative analysis we could not conduct is to assess the correlation between cell/embryo volume and the onset of bursty cell division events. This is true for both *C. elegans* and Zebrafish, and we have done a similar analysis using the C. elegans data comparing cell volume against developmental stage in [27]. Nevertheless, we can assume that sudden changes in the number of cells might reflect changes in embryo volume. This is particularly true in the Zebrafish example, as their embryos undergo numerous changes in shape over the developmental intervals of our analysis.

### Numeric Embryo Experiments

A numeric embryo (or perhaps more accurately a numeric one) allows us to understand the fundamental features of cell division events relative to the efficiency of their timing. Is one timing scheme superior to another? We know that in real (biological) lineage trees that cell divisions do not occur at a completely regular rate.

Are there advantages in one particular statistical signature over another, particularly when comparing it to an artificial (regular) scheme? Table 1 shows a summary of how this simulation is constructed.

**Table 1.** An example of our numeric simulation, with variable and sample values.

| Developmental Time Unit | Division Time (AU) | Generated Poisson Interval | Division Interval |
| --- | --- | --- | --- |
| 0 | 0 | 0 | 0 |
| 1 | 1 | 1 | 0 |
| 2 | 0 | 2 | 2 |
| 3 | 1 | 3 | 0 |

We use the uniform distribution as the basis for Poisson noise, which helps to execute things a bit faster on average. Compare this to uniform division times such as a division event occurring once every 20 units of time. Generated Poisson Interval represents the size of the interval between division events, while division Interval represents when the event occurs in developmental time. Our timing data can be modeled as branches of a binary tree which are generated every *n* units of developmental time. The intervals between *n*_*1*_, *n*_*2*_, *n*_*3*_,*…. n*_*t*_ are determined by a probability distribution, which can be uniform (every branching event occurring at completely regular intervals), or a Poisson distribution (where branching events are distributed in an exponential fashion).

The graphs in Figure 7 tells us that modeling division events using a Poisson distribution is that we can achieve the same number of divisions as fewer developmental time units. Figure 7 (top) shows a Uniform distribution of division events, while Figure 7 (bottom) shows the Uniform case as compared to other distributions (Exponential, Poisson, and Binomial). The Poisson distribution yields the “fastest” time relative to the number of divisions produced. By contrast, the Binomial distribution yields the lowest number of divisions (hence is the slowest method examined). However, none of these methods produce orders-of-magnitude differences in division rate, which is what would be expected from a bursty signature.

## Discussion

In this paper, we examine the periodicity of cell proliferation and division examined using three model systems: Zebrafish (*Danio rerio*), Nematode (*Caenorhabditis elegans*), and a simulated embryo. When we refer to periodicity in development, we mean events that reoccur over time. Regular pulses of cell proliferation events can occur in a short period of time. This leads us to propose a principle of development based on timing. There may also be a spatial component of developmental periodicity: these include signatures of time-independent spatial periodicity such as tilings and other repeatable patterns across space.

### Interpretation of Figures

We interpret Figures 1 and 3 in a number of ways. The first is by looking at components of variation over time. We measure this in terms of the interval between cell division times in *C. elegans* (Figure 2) and the frequency of cell division rates in Zebrafish (Figure 4). We also focus on intervals between other features in the time-series such as peaks for both species in Figure 5. In investigating peak intervals, we discover a similar distribution of cell division events between species in Figures 2 and 4, but a difference between species when looking at specific time-series features (Figure 5). The reason for this is clear: features such as peaks (magnitude) have a different underlying mechanism than events such as cell division. While both are linked to the lineage tree, magnitude differences are linked to the synchronization of cell division due to deterministic timing. With deterministic timing, synchronized cell divisions produce numerous cells at any one point in developmental time, but little fluctuation between time points. In the case of stochastic timing, several cells can be produced with a great degree of fluctuation between time points.

There are a number of ways to interpret the embryo network and 3-D plot shown in Figure 6. One interpretation is that in Zebrafish, the phenotype is built from the inside out, with densely-packed cells representing fledgling anatomical structures such as the notochord and heart. These clusters may be linked to rounds of cell division (occuring in temporal bursts), while cell divisions occurring during the inter-burst intervals may contribute to cells at the outer edge of the embryo and perhaps representing the ectoderm layer [28, 29]. In this way, temporal bursts of cell division lead to a spatial hierarchy of cell differentiation. Given an observed wave or peak in cell divisions at a certain point in developmental time, mitosis provides an opportunity to change gene expression, and ultimately serves as a collective signal for changes in cell fate. These phenomena demonstrate that developmental regulation is not simply a molecular mechanism, but a property of cell division and differentiation as well.

This spatial hierarchy involves a number of evolutionary and biophysical constraints that have been demonstrated in a number of experimental settings. For example, physical confinement affects the overall axial alignment and geometry of an embryo [30]. This includes our Zebrafish embryo network. Other types of fishes (*Astyanax*, see [31]) exhibit morphological changes in neural crest cell proliferation based on evolutionary changes due to ecological constraints. In *C. elegans*, asymmetrical cells (or daughter cells with significantly different volumes) result from physical constraints and compose 40% of *C. elegans* developmental cell divisions [32, 33]. Asymmetric cell divisions set up key cell-cell interactions [32] that are highlighted by the edges of embryo networks. Finally, by comparing nematic alignment of liquid crystals to spindles of mitotic cells, phase transitions in actively dividing cells are found to result from the timing of centrosome separation [33].

Figure 7 provides an introduction to the numeric embryo concept. In this Figure, we focus exclusively on the timing component of lineage trees. This is essentially a version of the time series shown for Zebrafish and *C. elegans* developmental time series, but with the temporal fluctuations smoothed out. These fluctuations are replaced with a cumulative sum of all cell division events occurring over a certain period of time. Comparisons between different distributions do not yield an appreciable difference in developmental speed (or the accumulation of *x* cells over a certain period of time). In Figure 7, all simulations were run for 60 iterations.

Investigating the potential of the Poisson distribution further, we investigate how this distribution approximates cumulative cell division (as was done in Figure 7) for three values of λ (0.1, 0.5, and 1.0). The outcome of this experiment is shown in Supplemental Figure 5. As this parameter value is increased, the number of cells per developmental time point increases while the interval between cell divisions decreases. While the function derived from λ = 0.1 is always slowest, the functions derived from λ = 0.5 and λ = 1.0 are similar for the first 20 timepoints, then diverge to reveal that λ = 1.0 clearly results in both faster cell divisions and a larger number of total cells after 200 iterations.

### Broader Questions

We can ask what it means when embryogenetic systems exhibit multiple pulses of cell proliferation from division events. In particular, the intervals between pulses provide information about the generative mechanisms behind production of the embryo. Our inquiry is particularly suited to quantitative interpretation, particularly in terms of characterizing “bursty” behaviors. These bursty behaviors are non-normally distributed generative processes [34] that describe the tempo and mode of development. While tempo and mode is generally an evolutionary phenomenon, these concepts also yield a model of developmental regulation that is explicitly temporal. Our result also suggests developmental regulation is not simply a molecular mechanism.

We can also speculate about the causes and breakdowns of observed periodicities (Table 2). Periodicities are caused by synchronized cell division and spatial proximities. The latter may be underlain by signaling mechanisms between neighboring cells, particularly in cases where cells are packed together along an axis or in a specific geometric configuration [35]. Periodicities are also caused by stochastic processes, which can result from many cells acting collectively without top-down coordination. Related to this is a phase transition, which results from a sudden, global change in embryo state. Phase transitions could be related to different stages of embryogenesis or other large-scale changes in cellular behavior. These periodicities can also break down due to similar factors such as desynchronization of cell division due to cell-cell signaling and interactions between the organism’s lineage tree and cell migration in the embryo. Stochastic processes and phase transitions can also break down over time. In the case of stochasticity, deterministic mechanisms can override the indeterminate behavior of cell (particularly cell division) and lead to more uniform divisions in time. In a like manner, phase transitions must break down (as they are quick and transformative), and in doing so enable the embryo to settle into a stable state.

**Table 2.** Causes and breakdowns of observed periodicities in embryos.

| **Causes** | **Breakdowns** |
| --- | --- |
| Synchronized Cell Division | Desynchronization Cell Division |
| Spatial Proximities | Lineage-Micratory Interactions |
| Stochastic Processes | Deterministic Processes |
| Phase Transition | Stability of State |

Our network analysis also demonstrates a connection between the spatiotemporal dynamics of cell division, cell differentiation, and systems-level view of timing. For example, we have found that structure and timing of interactions shape embryo network coherence signaling [36], which in turn is an indicator of diffusion between developmental cells that share network connections. While it is not discussed in this paper, gene expression fluctuations and stochastic noise in gene expression drives heterogeneity in division timing and even timing of differentiation [37, 38]. In particular, a focus on the molecular biology of the cell cycle across groups of developmental cells [39, 40] can provide more information about how fluctuations work in general at the single-cell level. Yet single cells acting in synchrony (or in the aggregate) define the patterns observed in our empirical data. One way to generalize our result to a broader cross-species context is to examine related phenomena such as mitotic bookmarking [41], in which heritable regulatory information is transmitted from mother to daughter cells in a cell lineage.

Our approach is also quite valuable [see 42, 43] for understanding this particular scale of the biological organism. For a fuller understanding in the context of groups of cells producing mean behaviors, we can appeal to the quantal mitosis hypothesis. Quantal mitosis involves changes in gene expression, in which the fate depends upon mitosis. This is also a gene expression-related memory mechanism that is widespread in development [41]. In cases of an observed wave or peak in cell divisions at a certain point in developmental time, mitosis provides an opportunity to change gene expression [44], and ultimately serves as a collective signal for changes in cell fate [45]. Finally, the way in which we decompose the spatiotemporal dynamics of the embryo might be useful as a supplement to reaction-diffusion models of morphogenesis [46]. Future work will involve extending this type of analysis to other species, in addition to developing our numerical models to include explicitly spatial phenomena.

## Supporting information

Supplemental Figure 1 full size

Supplemental Figure 2 full size

Supplemental Figure 3 full size

Supplemental Figure 4 full size

Supplemental Figure 5 full size

Supplemental Movie 1 link

## Acknowledgements

We would like to thank members of the DevoWorm group for their support and feedback, particularly Susan Crawford-Young. Thanks also go to the OpenWorm Foundation for their institutional support.

## Supplemental Movies and Figures

**Link:** https://commons.wikimedia.org/wiki/File:An-ensemble-averaged-cell-density-based-digital-model-of-zebrafish-embryo-development-derived-from-srep08601-s2.ogv

Supplemental Movie 1. A single-cell resolution time-series digital model of Zebrafish embryo based on light-sheet microscopy data from [7], licensed under Creative Commons Attribution 4.0 International.

**Supplemental Figure 1.**
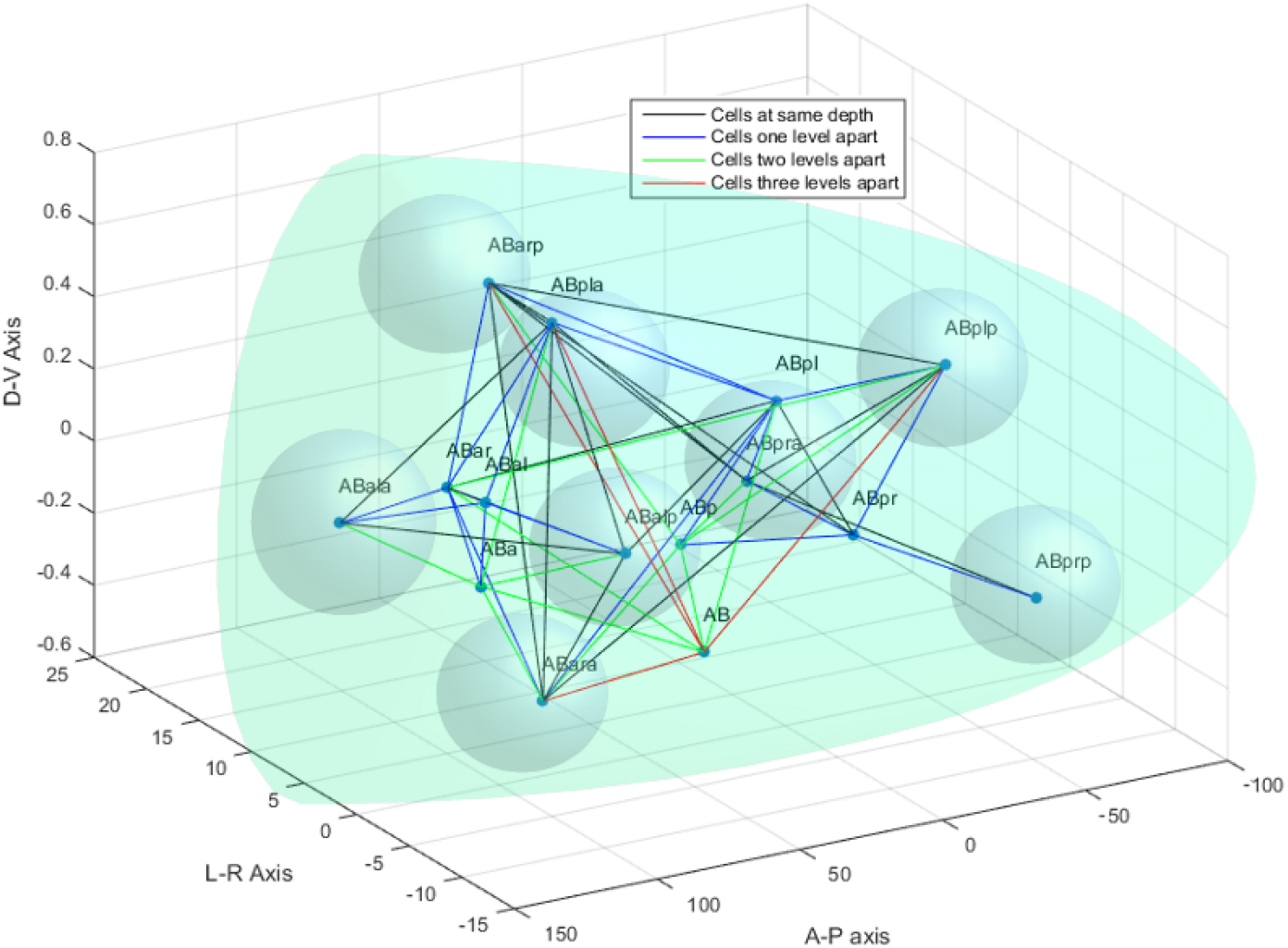
Example of an embryo network from the 16-cell *C. elegans* embryo build using cell tracking data. Data shown in the context of a cartoon showing the anterior end of the embryo. Different colored edges represent cells born at different generations of the lineage tree (levels).

**Supplemental Figure 2.**
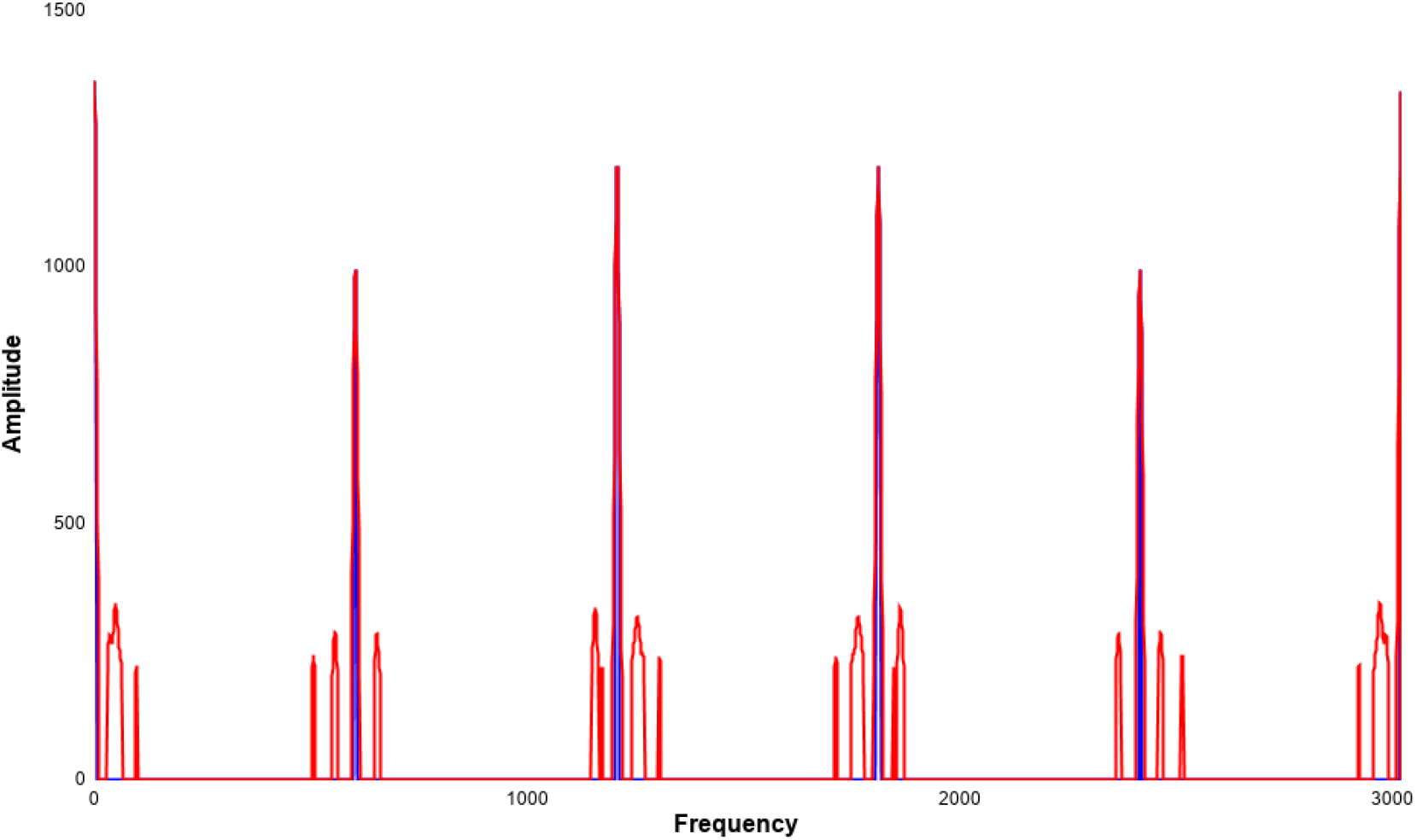
Frequency-domain plot of cell division event frequencies in *C. elegans* embryo. All events greater than an amplitude of 200 shown in red, while all events greater than an amplitude of 800 shown in blue.

**Supplemental Figure 3.**
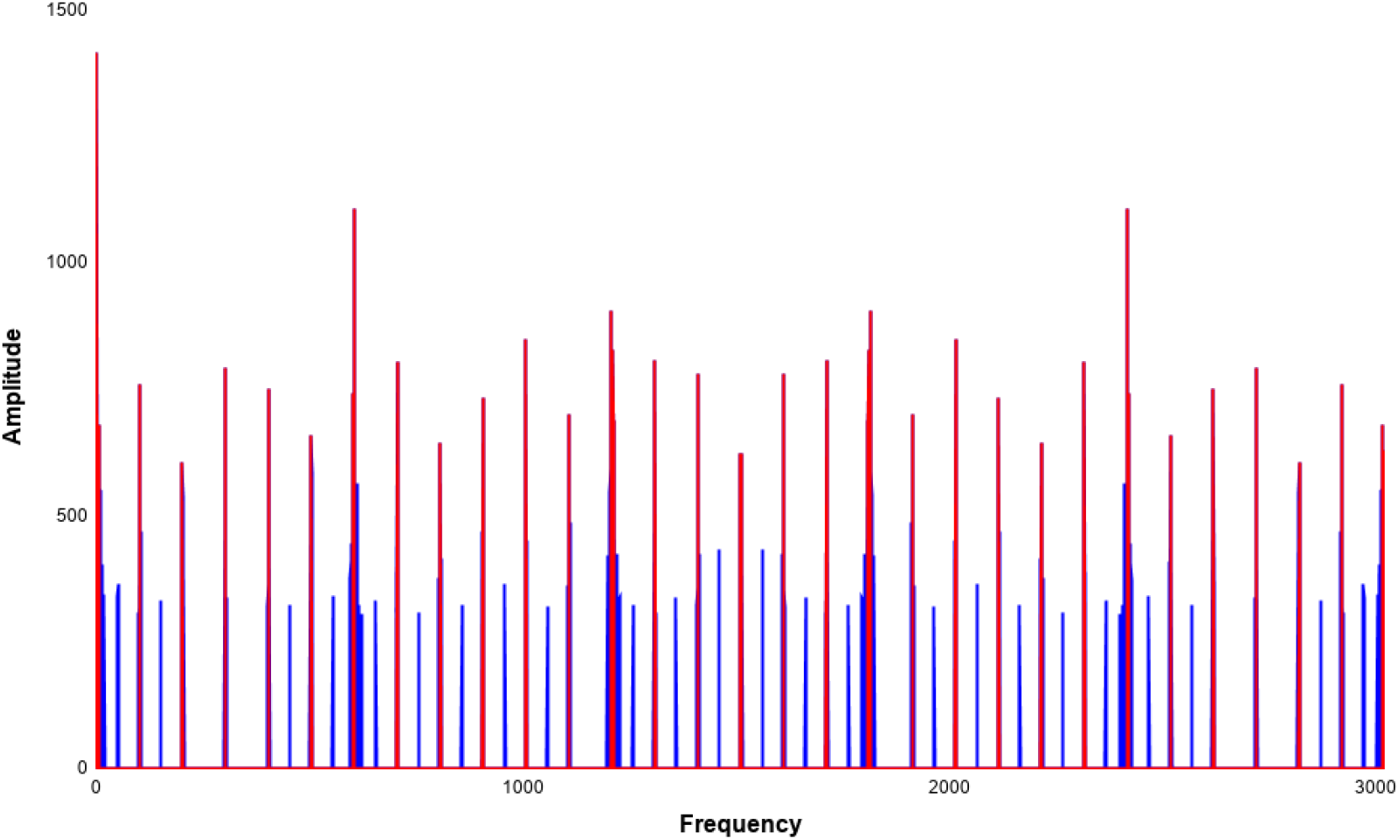
Frequency-domain plot of cell differentiation event frequencies in *C. elegans* embryo. All events greater than an amplitude of 300 shown in red, while all events greater than an amplitude of 600 shown in blue.

**Supplemental Figure 4.**
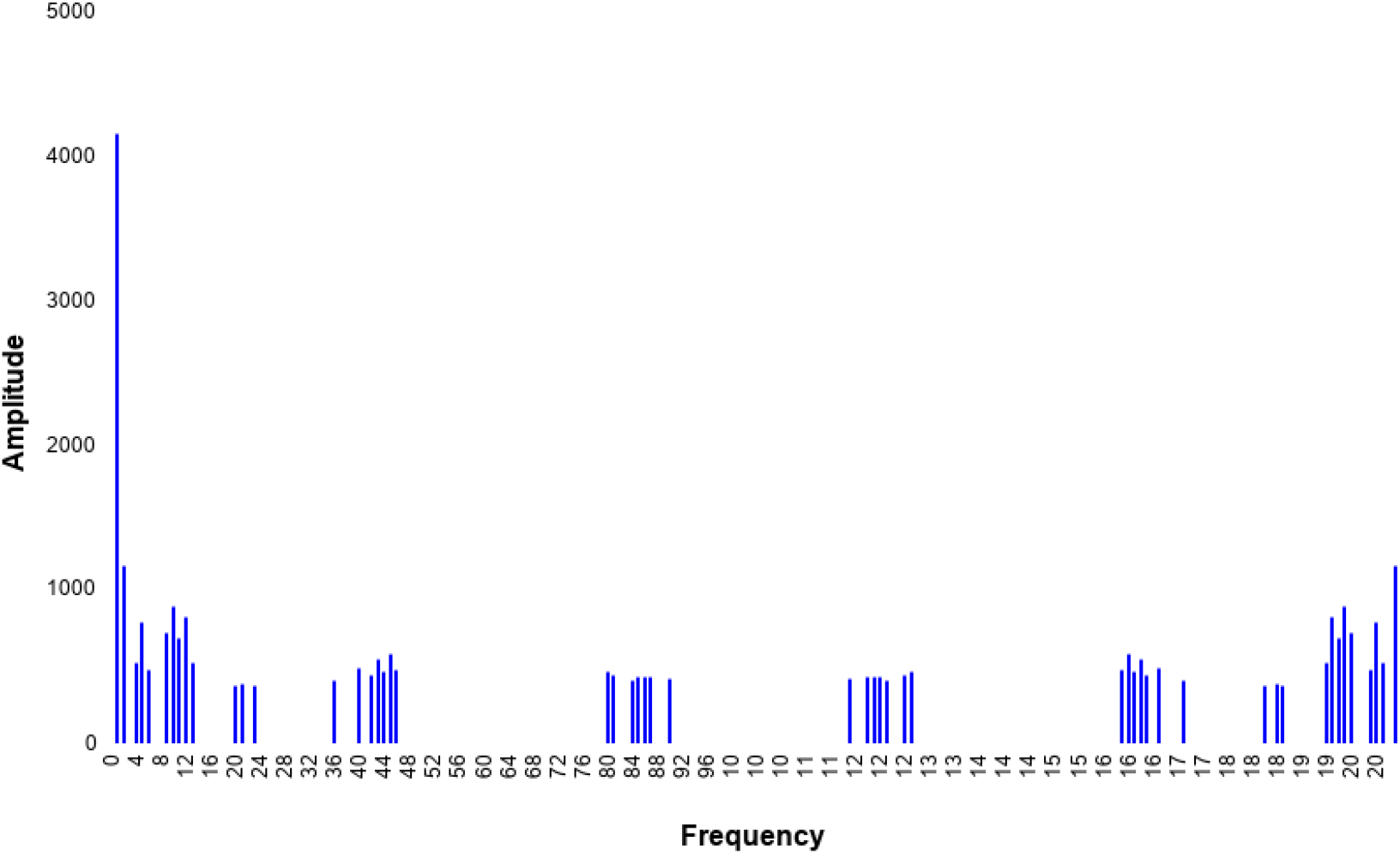
Frequency-domain plot of cell division event frequencies in Zebrafish embryo.

**Supplemental Figure 5.**
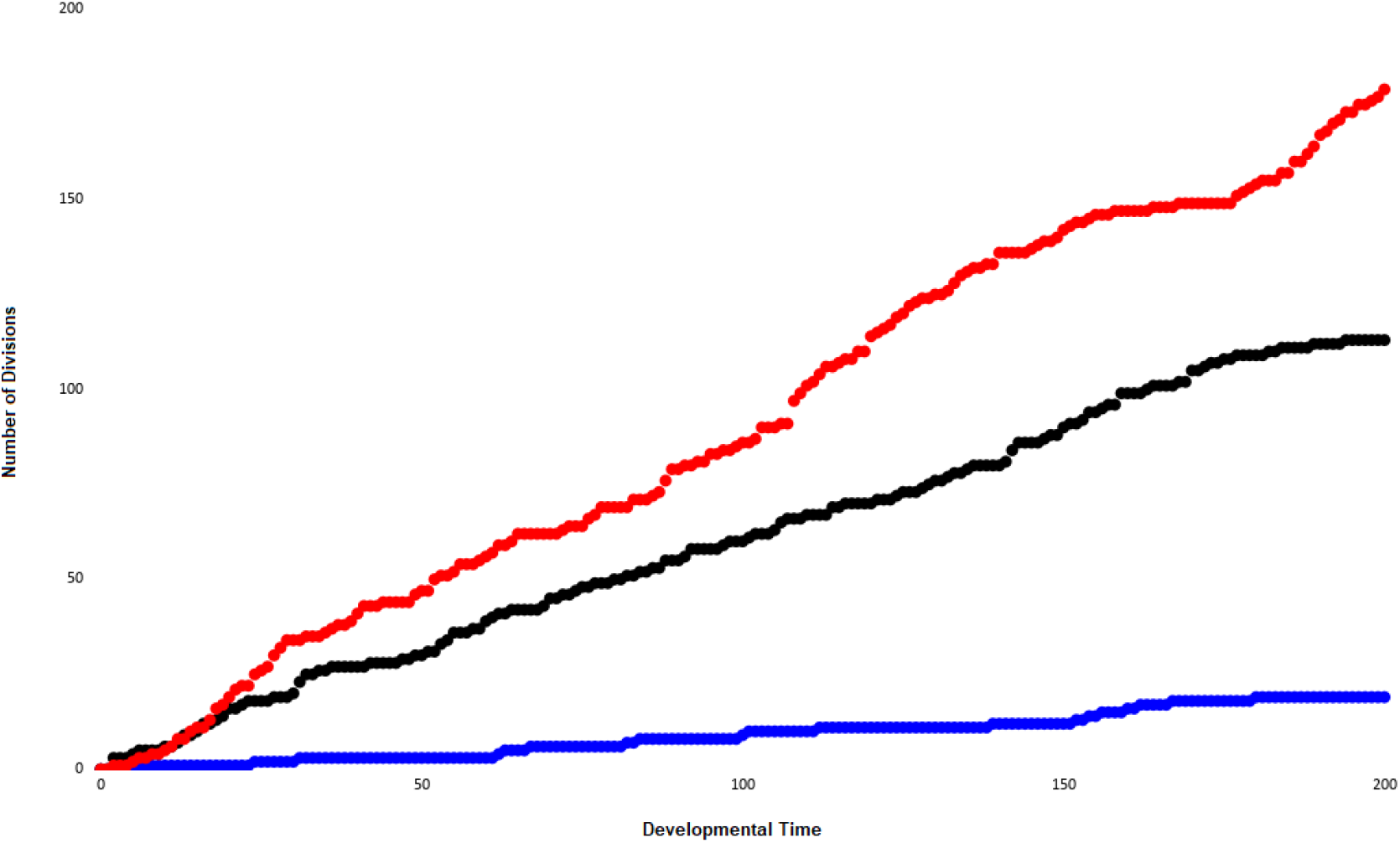
Comparison of cumulative cell division events and the speed of division generated by a numeric embryo for the Poisson distribution at three different values of λ. Blue: λ = 0.1, Black: λ = 0.5, Red: λ = 1.0.

