## Supplementary figures and images for "Periodicity in the embryo: emergence of order in space, diffusion of order in time"

### Supplemental Figure 1 full size

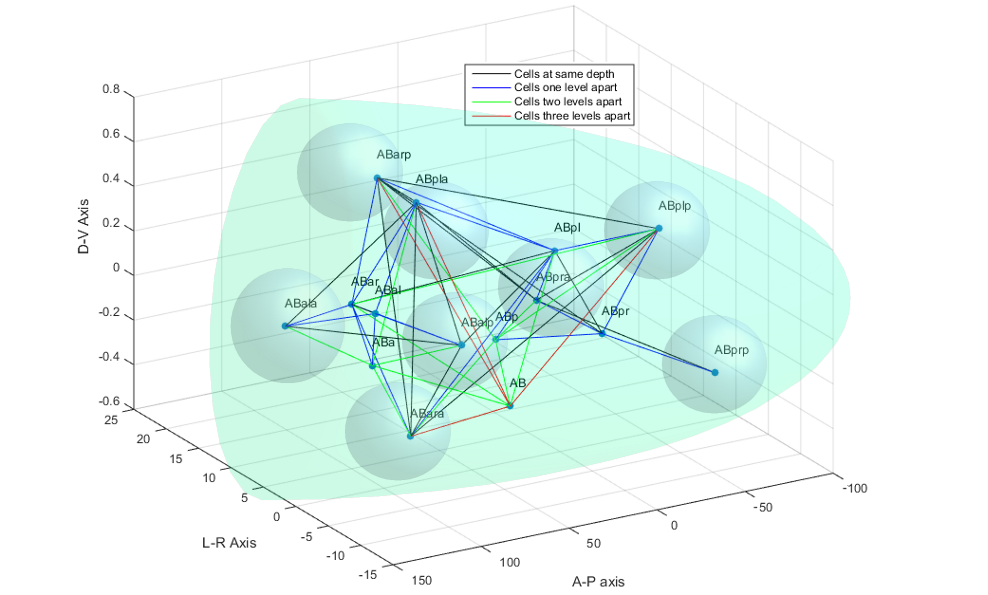

### Supplemental Figure 2 full size

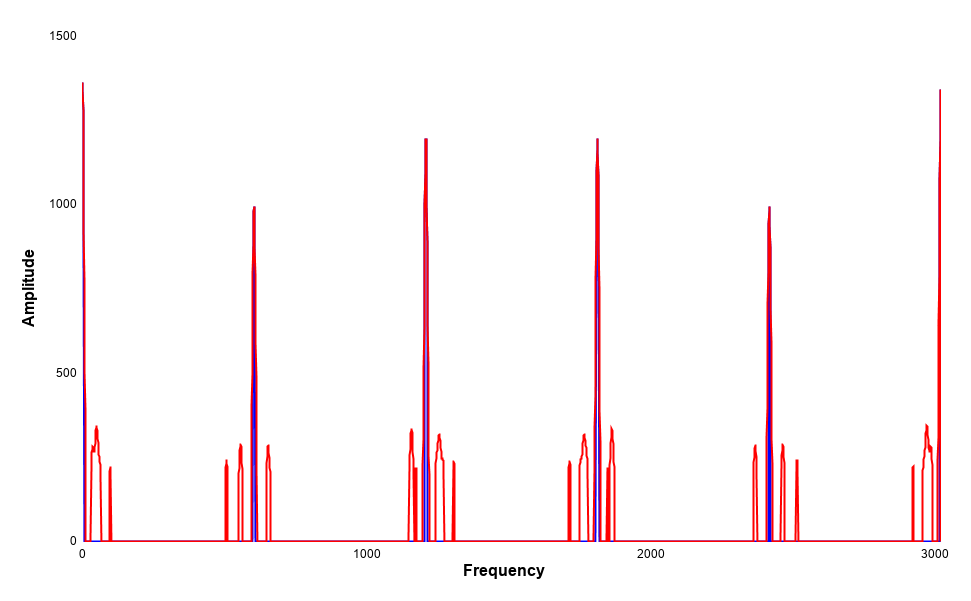

### Supplemental Figure 3 full size

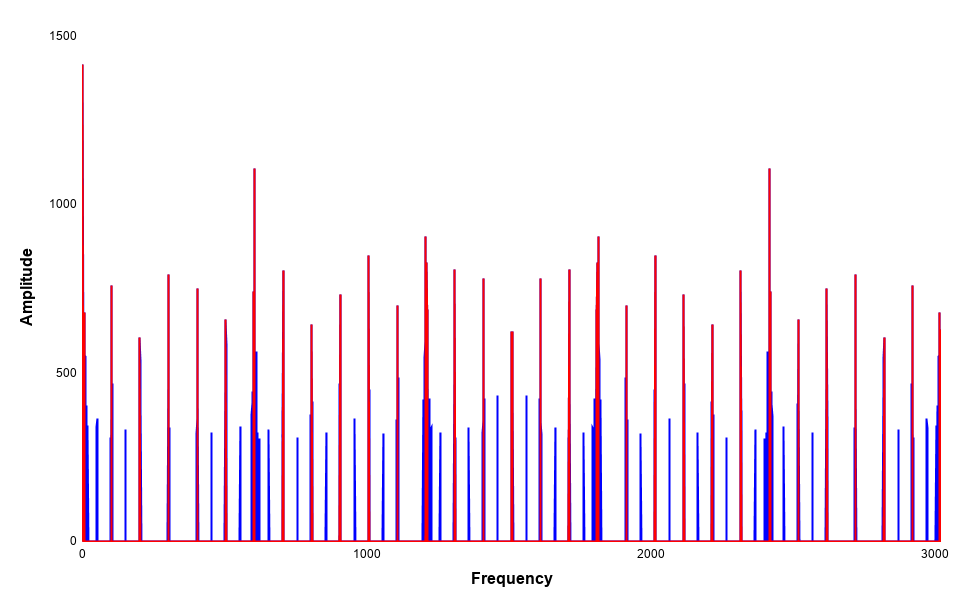

### Supplemental Figure 4 full size

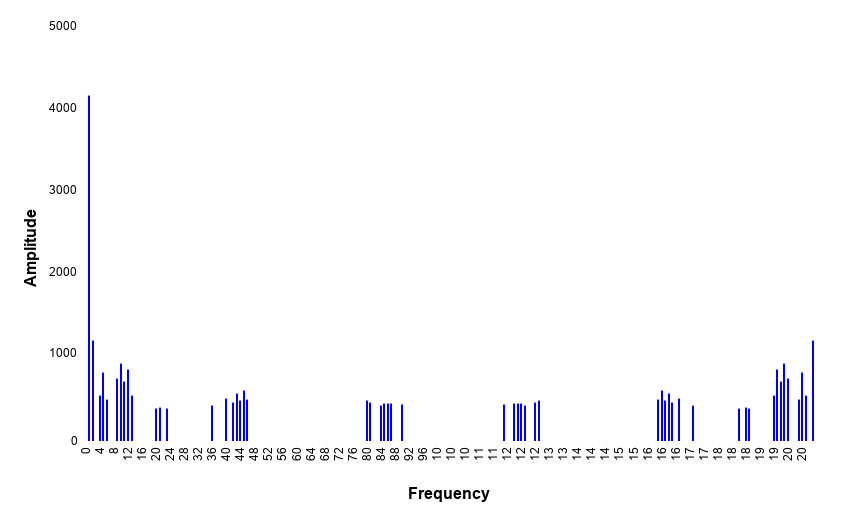

### Supplemental Figure 5 full size

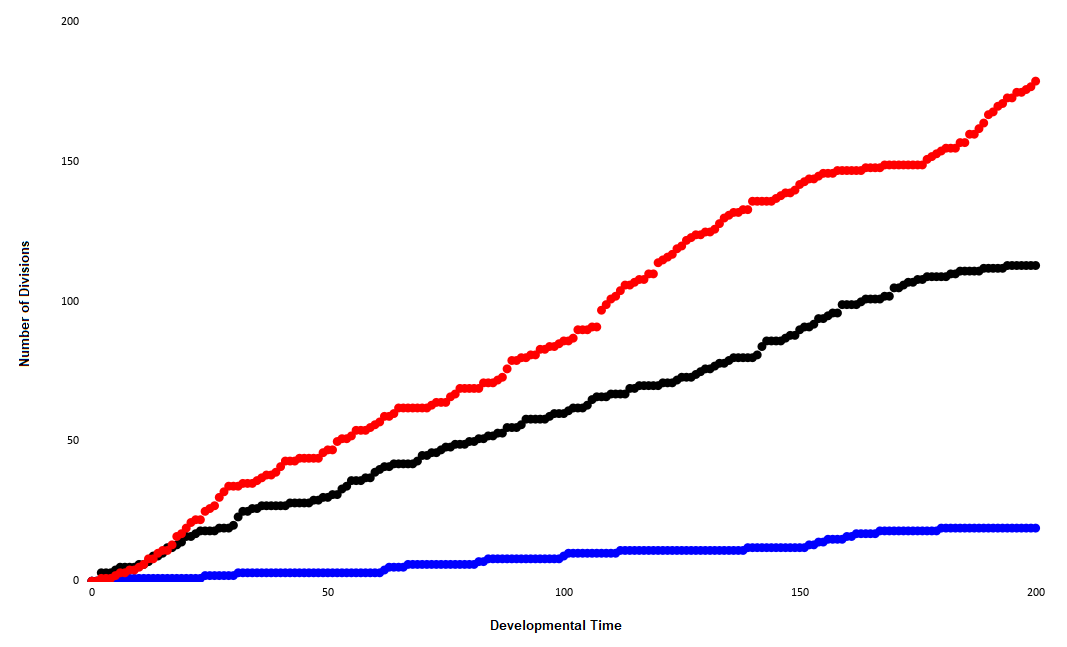
